# Hippocampal cells integrate past memory and present perception for the future

**DOI:** 10.1101/2020.03.08.982355

**Authors:** Cen Yang, Yuji Naya

**Affiliations:** School of Psychological and Cognitive Sciences, Peking University, Beijing, China; Academy for Advanced Interdisciplinary Studies, Peking University, Beijing, China; Center for Life Sciences, Peking University, Beijing, China; PKU-IDG/McGovern Institute for Brain Research, Peking University, Beijing, China; Beijing Key Laboratory of Behavior and Mental Health, Peking University, Beijing, China

## Abstract

The ability to use stored information in a highly flexible manner is a defining feature of the declarative memory system. However, the neuronal mechanisms underlying this flexibility are poorly understood. To address this question, we recorded single-unit activity from the hippocampus of two non-human primates performing a newly devised task requiring the monkeys to retrieve long-term item-location association memory and then use it flexibly in different circumstances. We found that hippocampal neurons signaled both mnemonic information representing the retrieved location and perceptual information representing the external circumstance. The two signals were combined at a single-neuron level to construct goal-directed information by three sequentially occurring neuronal operations (e.g., convergence, transference, targeting) in the hippocampus. Thus, flexible use of knowledge may be supported by the hippocampal constructive process linking memory and perception, which may fit the mnemonic information into the current situation to present manageable information for a subsequent action.

## Introduction

Declarative memory enables individuals to remember past experiences or knowledge and to use that information according to a current situation (1, 2). This flexible use of stored information is in contrast to procedural or fear-conditioned memory, in which acquired memory is expressed in a fixed form of associated actions or physiological responses (3-5). Previous studies revealed the involvement of the hippocampus (HPC) in the medial temporal lobe (MTL) in the formation and retrieval of declarative memory (2, 3, 6-14). However, the mechanism by which the HPC contributes to the flexibility in the usage of the declarative memory remains largely unknown.

The contribution of the HPC to declarative memory was often investigated by examining its spatial aspects in both human subjects (15-17) and animal models (3, 8-10, 13, 14, 18-20). In the preceding literature, the contributions of the HPC to the spatial memory task were successfully dissociated from those of the other brain areas when the start position in spatial mazes differed between training (e.g., “*south”* in a plus maze) and testing trials (e.g., “*north”*) (3, 21), because the fixed action patterns acquired during the training period (e.g., “*turn left*”) cannot guide the subjects to a goal position (e.g., “*west*”) in the testing trials. The HPC thus contributes to the memory task by retrieving a goal position, which could be represented in an acquired cognitive map (22). However, in order to reach the goal position, it is necessary but not enough for the subjects to remember the goal on the cognitive map. In addition, it would be critical for the subjects to locate their self-positions by perceiving current circumstances around them and relate the goal to the self-positions for a subsequent action. The subjects thus need to transform the goal position within the cognitive map into goal-directed information relative to a specific circumstance (i.e., start position) that the subjects currently experience. In the present study, we hypothesized that the goal-directed information in the specific circumstance may be constructed by combining the retrieved memory and incoming perceptual information on the same principle as “*constructive episodic memory system*” (1, 23). In their theory, the constructive episodic memory system recombines distributed memory elements for both remembering the past and imagining the future (1, 24). However, the neuronal operations for the “*mental time travel*” (25) to both the past and future have not been identified at the single-neuron level.

We therefore investigated whether and how the HPC neurons combine the retrieved location with the perceived circumstance in order to construct goal-directed information. To achieve this purpose, we devised a new memory task for macaque monkeys, in which memory retrieval and its usage were separated by sequential presentations of two cues in a single trial (Fig 1). The first cue presented a visual item (item-cue) that would trigger retrieval of the location associated with the item. The second cue presented a background image (background-cue) that would be combined with the retrieved location to construct goal-directed information. This task structure allowed us to separate the constructive process from the retrieval of item-location association memory. We referred to this new task as the constructive memory-perception (CMP) task. We found a constructive process operated by the HPC neurons, which started from a “*convergence*” process that combines the retrieved memory triggered by the first cue and the incoming perceptual information presented by the second cue. Then, the HPC neurons transiently exhibited the “*transference*” process from the retrieved location into a target location, and the HPC finally represented the target location itself (“*targeting*” process). These sequential neuronal operations thus provide a goal-directed signal by fitting the retrieved memory into the current situation. These findings suggest that the HPC equips the declarative memory with flexibility in its usage by the constructive process combining memory and perception.

**Fig 1.**
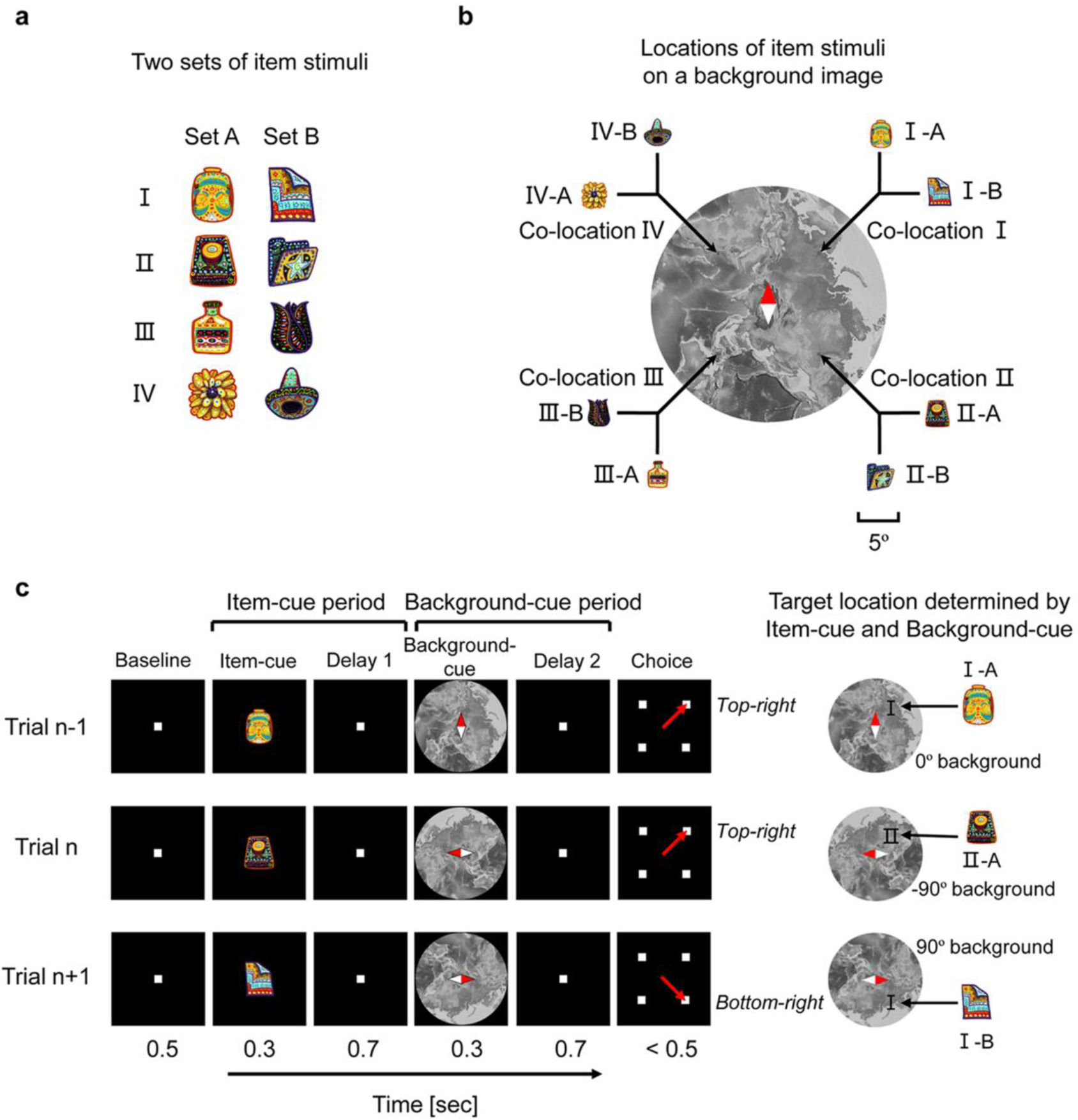
Constructive memory-perception (CMP) task. **(a)** Item stimuli. **(b)** Item-location association pattern. Two items, one from set A (e.g., I-A) and the other from set B (e.g., I-B), were assigned to each location (e.g., co-location I) on the background image. Scale bar for both item-cue and background-cue stimuli, 5° visual angle. **(c)** Schematic diagram of the CMP task. An item-cue and background-cue were chosen pseudorandomly in each trial. The monkeys should maintain fixation on the center until the end of the background-cue period including Delay 2, then saccade to the target location (red dashed arrow) during the choice period. Correct target location determined by both the item- and background-cues (the assigned location of item- cue on the current background-cue). Monkeys were trained using every 0.1° step in orientation from −90° to 90°, though only five orientations (−90°, −45°, 0°, 45° and 90°) were tested during the data acquisition. Relative sizes of the item-cue stimuli to the background-cue stimuli were magnified for display purpose.

## Results

### Constructive memory-perception (CMP) task

Two rhesus macaques were trained to perform the CMP task. In the CMP task, four pairs of visual items were assigned to four different locations (*co-locations*) on a background image (Figs 1a and 1b, S1 Text). We referred to the two items in each pair as “*co-location*” items (e.g., I-A and I-B) because the two items were assigned to the same location on the background image. The configuration of the co-location items allowed us to evaluate an item-location memory effect for each single neuron by examining the correlation in its responses to the co-location items. In the present study, we used the same eight visual items and one background image during all the recording sessions. In each trial, one of the eight items was presented as an item-cue (e.g., II-A) (Fig 1c). After a short delay, a randomly oriented background was presented as a background- cue (e.g., −90°). The subjects were then required to saccade to the target location (e.g., top-right position on the display) determined by the combination of the item- and background-cues (e.g., co-location II on the −90°-oriented background on the display).

In the initial training, the monkeys learned the item-location association through trials with a fixed orientation of background-cue (that we defined as 0°). After they learned the association between items and locations in trials with the 0° background-cue, orientation of the background- cue was randomly chosen from −90° to 90° (in 0.1° steps). During the recording session, the orientation of the background-cue was pseudorandomly chosen from among five orientations (− 90°, −45°, 0°, 45°, and 90°). The necessity to choose the associated location on the oriented- background required the monkeys to associate the visual items with locations relative to the background rather than associate the visual items with motor responses directly. Moreover, the temporal separation of the two cue presentations allowed us to trace the transformation process from the retrieved information to the goal-directed information in the CMP task. While the monkeys performed the task, we recorded single-unit activity from 456 neurons in the HPC of the MTL (S1 and S2 Figs and S1 Table).

### Representation of the retrieved memory

We first investigated the retrieval process during the item-cue period of the task. Figure 2a shows an example of a neuron exhibiting item-selective activity (item-selective neuron, *P* < 0.01, one-way ANOVA). This neuron exhibited the strongest response to item I-A (*optimal*), while an item paired with the optimal item I-B (*pair*) elicited the second-strongest response from the same neuron (Fig 2b). The neuron thus strongly responded to only the particular co-location items (i.e., I-A and I-B) but not to others. The selective responses to I-A and I-B could not be explained by eye position (S3 Fig). These results may suggest that the neuron represented the co-location that was associated with the items I-A and I-B. We examined the association effect by calculating the Pearson correlation coefficient between the responses to the four pairs of co-location items and referred to it as the co-location index. The co-location index of this neuron was significantly positive (Fig 2b) (*P* < 0.0005, permutation test, one-tailed), which suggests that the item-selective activity of this neuron signaled the co-locations retrieved from item-cues. Figure 2c shows the population-averaged spike density functions (SDFs) of item-selective neurons (n = 136) to optimal items, their pairs, and other items (average across six items). The responses to the items paired with the optimal items were significantly larger than those to the other items during the item-cue period (*P* < 0.01 for each time step, *t*-test, two-tailed). We also calculated the co-location index for the item-selective neurons. Their indices showed extremely large values (*r* = 0.89, median) (Fig 2d), indicating that item-selective activities in the HPC reflected the co-location information assigned to the individual items. We confirmed that the co-location effect on the item-selective activities could not be explained by eye position (S2 Text). These results suggest that the HPC exhibited strong location signals retrieved from item-cues.

**Fig 2.**
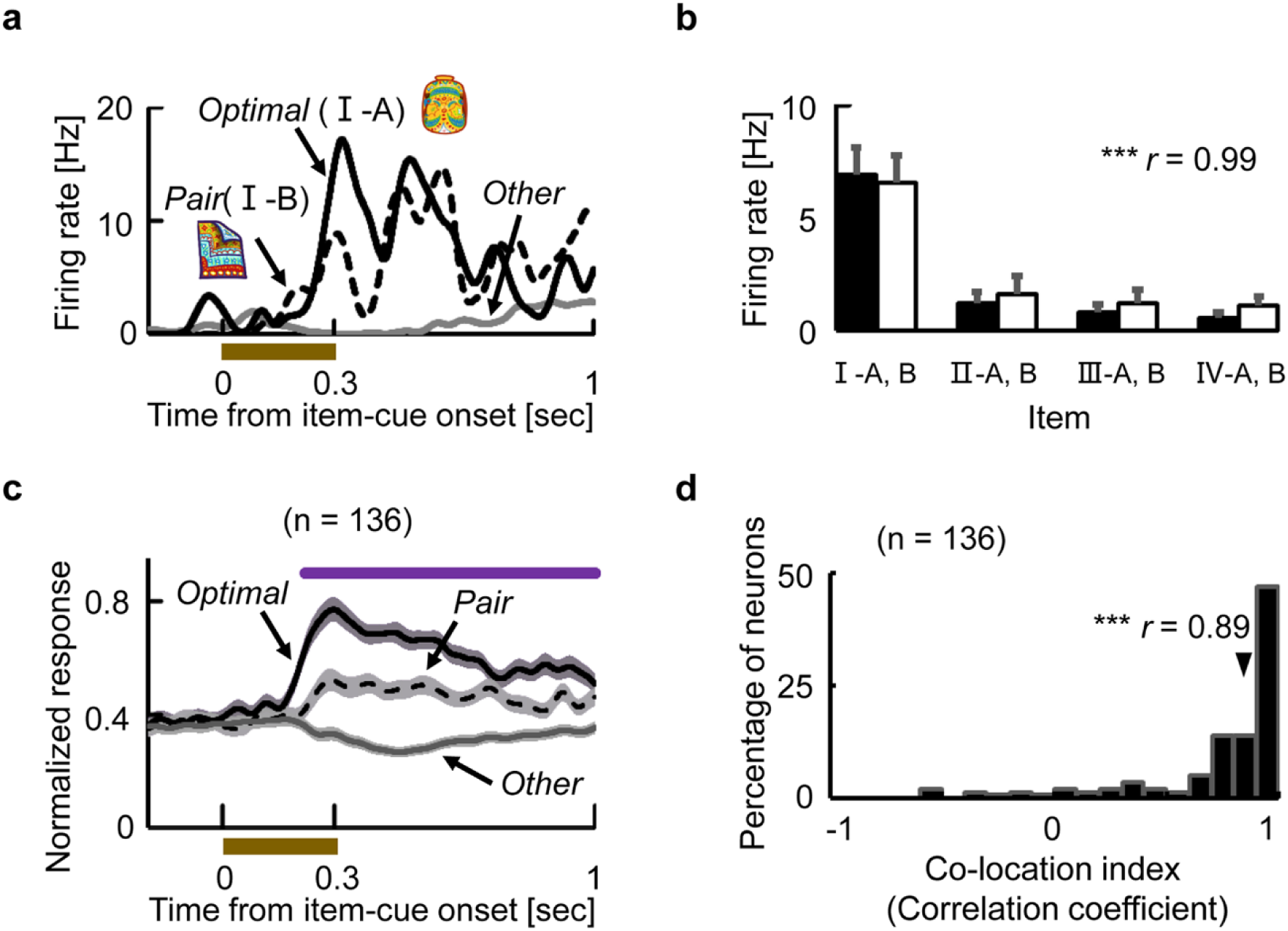
Representation of the retrieved memory. **(a)** Example of an item-selective neuron. Black lines indicate SDFs in trials with the best co-location items (i.e., *Optimal* and *Pair*). Grey line indicates averaged response in the other trials (*Other*). Brown bar indicates presentation of the item-cue. **(b)** Mean discharge rates and SEM of the same neuron in Figure 2a during the item-cue period for each item. Black bars, set A. White bars, set B. *r*, co-location index. *** *P* < 0.0005, Permutation test, one-tailed. **(c)** Population-averaged response of item-selective neurons (n = 136). SDFs in trials with the best co-location items (i.e., *Optimal* and *Pair*) and other items. Shading, SEM. Purple line, time duration indicating a significant (*P* < 0.01, t-test, two-tailed) difference between *pair* and *other*. **(d)** Distributions of co-location indices for item-selective neurons (n = 136). *r*, median value. *** *P* < 0.0001, Wilcoxon’s signed-rank test.

### Convergence of the retrieved memory and incoming perception

We next investigated how the incoming background-cue information affected the retrieved location signal. Figure 3a shows an example of a neuron exhibiting selective responses to the background-cues (*P* < 0.01, three-way nested ANOVA, see also S3 Text). This neuron showed the strongest responses across all the co-locations when the orientation of the background-cue was 90°, while it showed only negligible responses when the orientation was −90°. In addition to the neurons showing only background-selective activity (e.g., Fig 3a), we found neurons showing selectivity for both co-locations and backgrounds. An example neuron in Figure 3b began signaling co-locations III and IV at the end of the item-cue period. After the background- cue presentation, this neuron exhibited additional excitatory responses for the preferred co-locations (i.e., III and IV), especially when the orientation of background-cue was 90°. The background-selective responses were thus combined with the co-location-selective responses in this individual neuron (see also S4a Fig). We evaluated the similarity of orientation tuning across co-locations for the example neuron by calculating the Pearson correlation coefficient between the responses to the different orientations of the background-cues for the “best co-location” (III) and those for the “second-best co-location” (IV), and found high similarity of orientation tuning across the co-locations (*r* = 0.95). These results indicate that this neuron received the same background-cue signal irrespective of which co-location signal the neuron held from the item-cue period.

**Fig 3.**
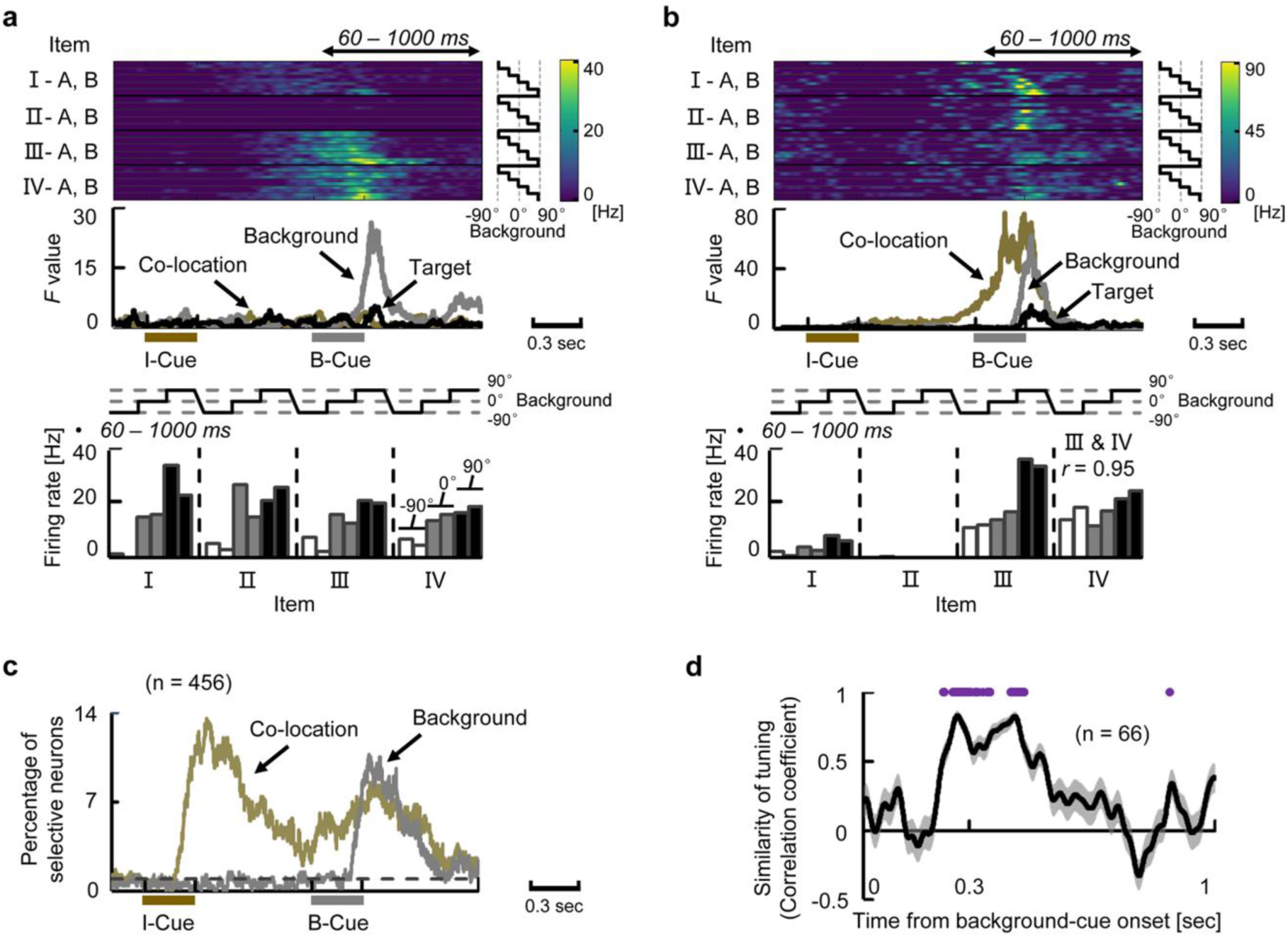
Convergence of the retrieved memory and incoming perception. **(a)** Example neuron signaling background effect. (Top) Each row contains an SDF for each combination of item- and background-cues. (Middle) Time courses of *F* values. Brown bar, presentation of the item-cue. Grey bar, presentation of the background-cue. (Bottom) Mean discharge rates for each combination of item- and background-cues during 60-1000 ms after background-cue onset. White, grey, and black bars indicate −90°, 0°, and 90° orientations of background-cue, respectively. Two bars with the same color indicate the co-location items (e.g., left bar, I-A; right bar, I-B). **(b)** Example neuron signaling background and co-location effects in a “*convergent*” manner. Same format as Figure 3a. *r*, Pearson correlation coefficient between the orientation tunings (responses to −90°, 0°, and 90°) for “best co-location” (III) and for “second-best co-location” (IV). **(c)** Time courses of percentages of co-location-selective and background- selective neurons (recorded neurons, n = 456). Dashed line, chance level = 1%. **(d)** Time course of similarity of orientation tuning. Line and shading, means and SEMs of the similarity of the orientation tunings for co-location-selective neurons. Purple line, time duration in which the similarity was significantly positive (*P* < 0.01, n = 66, Wilcoxon’s signed-rank test for each time step).

After background-cue presentation, a substantial number of neurons (22% of the recorded neurons, *P* < 0.01, three-way nested ANOVA) exhibited either co-location-selective activities (14%) or background-selective activities (14%). Interestingly, the proportion of co-location-selective neurons increased after the background-cue onset (Fig 3c). We evaluated the background-cue effect on the co-location-selective activities in the HPC by examining the similarity of orientation tuning for each of the co-location-selective neurons (n = 66). A similarity between the orientation tunings was observed from 228 to 458 ms after the background-cue onset in the population (*P* < 0.01 for each time step, Wilcoxon’s signed-rank test) (Fig 3d). These results showed that the background-cue information signaling its orientation converged on the co-location signal retrieved from the item-cues at the single-neuron level in the HPC (“*convergent-type*”), which indicates that the HPC combined the memory and perceptual signals.

### Representation of retrieved location and target location

After the background-cue presentation, some co-location-selective neurons started to exhibit target location selectivity. For example, a neuron in Figure 4a responded to item-cues that were assigned to the co-location III during the item-cue period, while the same neuron showed selective activity for a particular target location that corresponded to the bottom-left on the display (*yellow*) during the background-cue period. The bottom-left of the target location matched to the co-location III if we assume the background image with 0° orientation (Fig 4b). The responses to the target locations during the background-cue period were largely correlated with those to the co-locations during the item-cue period when the co-locations were assumedly positioned relative to the 0° background image (matching index, *r* = 0.99), but not to the −90° (*r* = −0.28) nor 90° (*r* = −0.23) background image (Fig 4b). This result may imply that the item-location is retrieved relative to the 0° background image as default. The presence of the default position/orientation of the background image in a mental space of the monkeys may reflect the effect of initial training, during which the monkeys learned the combinations of items and locations in trials with the 0° background-cue. To test this implication, we first collected 49 neurons showing significant target selectivity out of the 136 neurons with item selectivity during the item-cue period. These neurons tended to show the preferred target locations that corresponded to the preferred co-locations relative to the 0° background-cue (default orientation), but not to the other orientations (Fig 4c). These results support the presence of the default position/orientation of the background image for the representation of the retrieved item-location in the HPC. This implies that at least some HPC neurons may represent the retrieved location, which can be linked to a particular position in a real space rather than a conceptual category classified by the co-locations.

**Fig 4.**
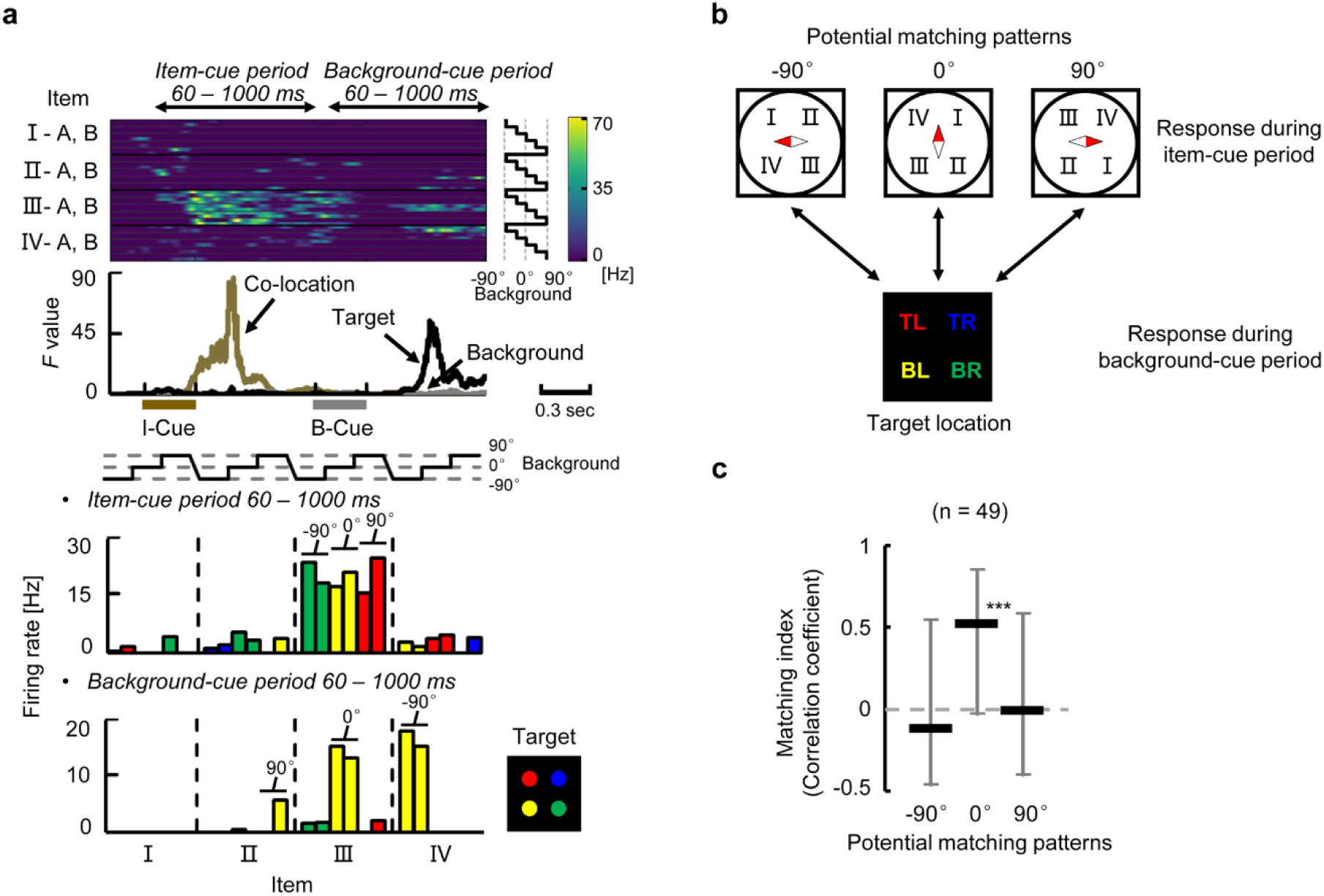
Representation of retrieved location and target location. **(a)** Example neuron signaling a particular co-location during the item-cue period and a particular “*targeting*” location during the background-cue period. Same format as Figure 3a, except that target locations in the bar graph are indicated by colors (bottom panel). For example, yellow color corresponds to the bottom-left of the target location on the display. **(b)** Potential matching patterns between the co-location and target location. I-IV, co-location I-IV. TR, top-right; BR, bottom-right; BL, bottom-left; TL, top-left. **(c)** Matching index for each matching pattern (using neurons signaling both co-location-selectivity during the item-cue period and target-selectivity during the background-cue period, n = 49). Black bar, median value. Error bar, quarter value. *** *P* < 0.0001, Wilcoxon signed-rank test.

### Construction of the goal-directed information

We next investigated how the retrieved location was transformed to the target location when the background-cue was presented. Figure 5a shows an example neuron that exhibited strong transient responses to particular combinations of item-cues and background-cues [(I, 90°) and (III, −90°)] corresponding to the same target location (bottom-right on the display, *green*). However, this neuron did not show the transient responses when the background-cue was 0° even though the combination (II, 0°) corresponded to the same target location (bottom-right). This implies that the neuron responded only when the retrieved location was transferred to the preferred target location (i.e., bottom-right). This “*transferring-type*” of activity contrasts to the target selective activity of the neuron shown in Figure 4a, which signaled the preferred target location itself regardless of the co-locations of item-cues or the orientations of background-cues. We refer to the latter type of target-selective activity as “*targeting-type*” (Fig 4a, S4b Fig). Interestingly, some individual neurons exhibited *convergent-type* activity first, then *transferring-type* activity, and finally *targeting-type* activity (Figs 5b, S4c Fig). These results imply a temporal relationship between the *transference* effect and *targeting* effect during the construction of goal-directed information.

**Fig 5.**
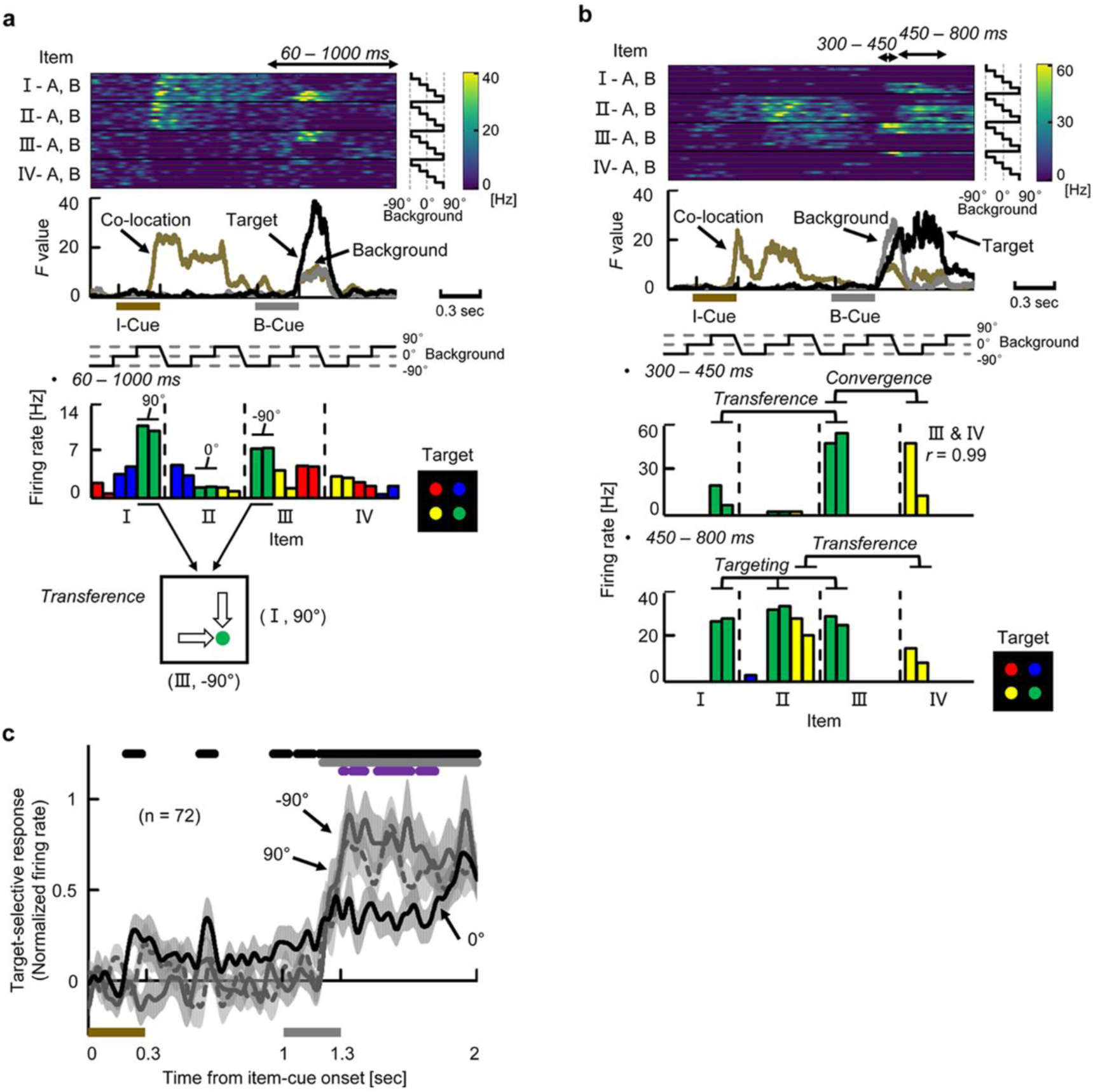
Construction of goal-directed information. **(a)** Example neuron showing the “*transference*” effect (I & 90° and III & −90° for the bottom-right). Same format as Figure 4a. Bottom panel shows a schematic diagram of “*transference*” from the retrieved location (co-locations I and III) into the same target location (bottom-right, *green*). **(b)** Example neuron showing multiple operations. **(c)** Target-selective responses (“best” minus “others”) in trials with −90°, 0°, and 90° background-cues for target-selective neurons (n = 72). Curves and shading depict means and SEMs of population-averaged SDFs. Top lines, time duration in which target-selective responses (“best” minus “others”) were significantly positive in trials with 0° background-cues (*black*) and with either −90° or 90° background-cues (*grey*) (*P* < 0.05, t-test, two-tailed). Purple line, time duration in which the “best” target responses were significantly larger in trials with either −90° or 90° compared with 0° background-cue (*P* < 0.05).

To examine the temporal relationship at the population level in the HPC, we compared time courses of the two types of target-related effects for the target-selective neurons (n = 72) by examining the effect of background-cues in different orientations (−90°, 0°, and 90°) on the target-selective responses (Fig 5c). The target-selective responses in trials with the −90° and 90° background-cues became significantly larger than those with the 0° background-cue from 309 to 786 ms after the background-cue onset (Fig 5c). The increase in target-selective responses after the −90° and 90° background-cues may reflect the transfer of the retrieved location into the preferred locations of individual HPC target-selective neurons (“*transferring-type*”). Then, the target-selective responses in trials with the 0° background-cue began to increase in the middle of delay 2, and the target-selective responses ultimately became indistinguishable among all the background-cues (Fig 5c), which may represent the target locations themselves (“*targeting- type*”). In trials with a 0° background-cue, target-selective responses were observed not only during the background-cue period but also during the item-cue period (*P* < 0.05, t-test, two-tailed) (Fig 5c), which confirmed the presence of the default position/orientation of the background image for the representation of the retrieved item-location in the HPC. Considering the fact that the immediate background-cue effect converged on the retrieved location signal, these results suggest involvements of sequential neuronal operations of *convergence, transference*, and *targeting* in the constructive process in which both memory and perception were combined to generate a goal-directed representation of the memory.

### Neuronal signal predicts animals’ behavior

We finally investigated whether the target-selective responses in the HPC were correlated with subjects’ behaviors. For this purpose, we conducted an error analysis for the target-selective activities during the background-cue period. Figure 6a shows an example neuron exhibiting target-selective activities. This neuron showed strong responses during the background-cue period when the animal chose the top-left position (red) not only in the correct trials (Correct trials, red) but also the error trials (False Alarm, black). In contrast, the neuron did not respond when the animal made mistakes by missing the top-left target position (Miss, grey). We examined whether the target-selective activities in the error trials could be explained by the positions animals chose or the correct positions of the trials using partial correlation coefficients (see Methods). The activities in the error trials of this neuron were related with the animals’ choice (*r* = 0.51, *P* < 0.0001) but not with the correct position (*r* = −0.18, *P* = 0.94). We calculated the partial correlation coefficients for the target-selective neurons with more than 10 error trials and found that the activities in error trials reflected the animals’ choice rather than the correct position (*P* < 0.0005, Wilcoxon’s signed-rank test) (Fig 6b). These results suggest that the target-selective activity constructed by the HPC neurons predicts the subsequent animal behavior.

**Fig 6.**
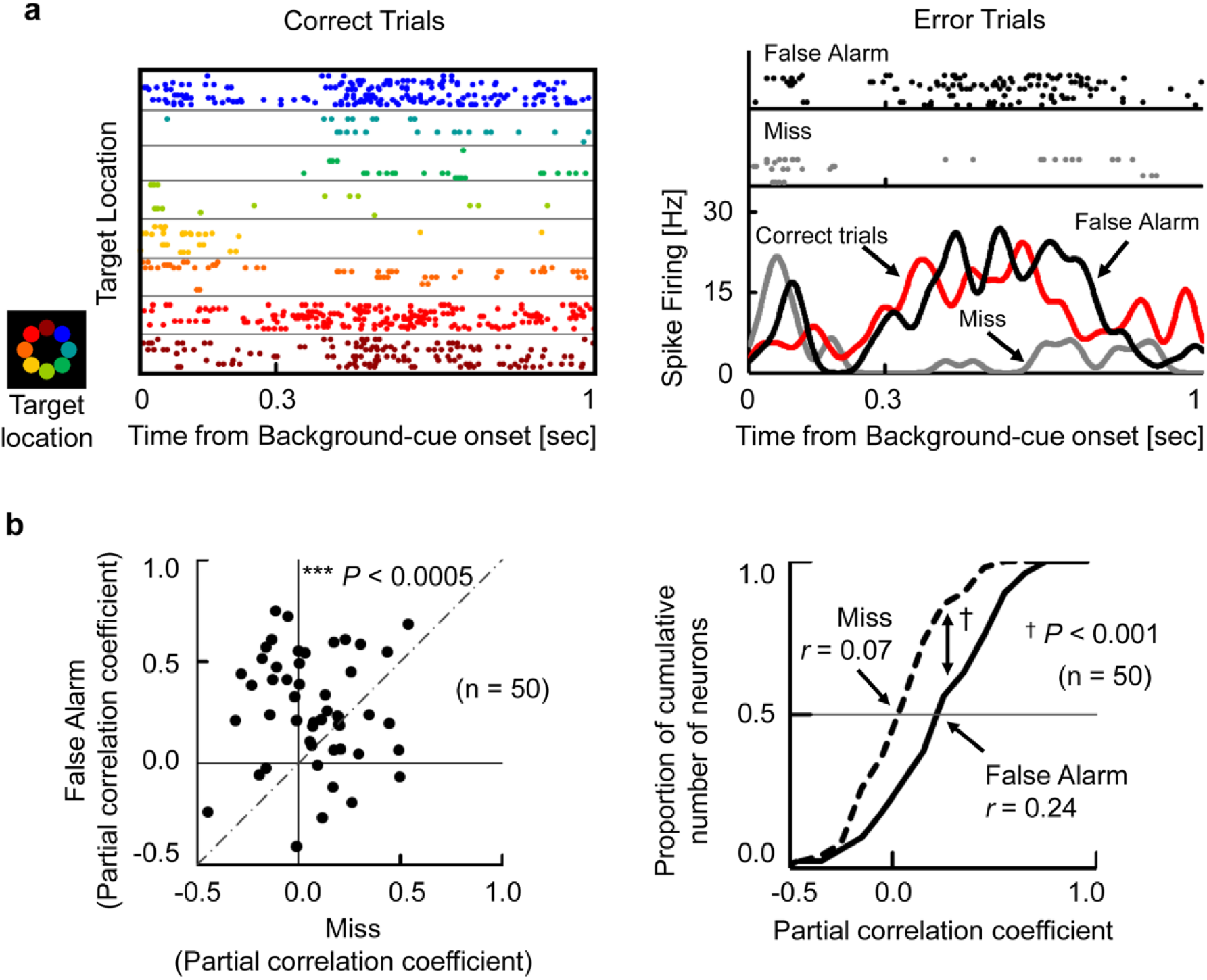
Neuronal signal predicts animals’ behavior. **(a)** Example neuron exhibiting target-selective activity. (Left) Raster displays of correct trials sorted by target locations. Colors indicate target locations on display. The neuron exhibited preferred responses when the target location was the top-left (*red*). (Right) Raster displays of error trials and SDFs (σ = 20 ms). False Alarm, top-left as the incorrect positions the subjects chose (*black*). Miss, top-left as the correct positions the subjects missed (*grey*). Correct trials, top-left as the correct positions the subjects chose (*red*). **(b)** Error analysis for target-selective neurons with at least 10 error trials (n = 50). False Alarm, the false positions the subjects chose. Miss, the correct positions the subjects missed. (Left) Scatter plot of partial correlation coefficients for individual neurons. Each dot indicate one neuron. *** *P* < 0.0005, Wilcoxon’s signed-rank test. (Right) Cumulative frequency histograms of partial correlation coefficients. *r*, median value. † *P* < 0.001, Kolmogorov- Smirnov test.

## Discussion

The present study aimed to investigate whether the flexible use of past knowledge can be explained by a constructive process in the HPC. We found a robust memory signal reflecting the location information retrieved from an item-cue (Fig 2, S5 Fig), which was even enhanced after presentation of the background-cue (Fig 3c). The perceptual information of the background-cue was converged on the retrieved location signal (Fig 3, S5 Fig), and then the retrieved location was transferred to the target location assigned by the combination of the two cues in each trial (Figs 5, S5 Fig). The target information signaled by the HPC neurons was correlated with the animal’s subsequent behavioral response (Fig 6). The present findings thus indicate that the HPC neurons combine mnemonic information with perceptual information to construct goal-directed representations of the retrieved memory, which would be useful for a subsequent action in the current situation.

In previous electrophysiological studies, the effects of the item-location association memory were presented as learning-dependent changes in firing rates (e.g., changing cells) (8, 26) or selective responses to a particular combination of items and locations (26-28). However, these studies did not identify the location signal retrieved from an item-cue. In the present study, we evaluated the item-location association memory as correlated responses to the co-location items, which may reflect the retrieved location (Fig 1a). The correlated responses to semantically linked items were previously investigated using the “pair-association” task (29-31). Different from the co-location items linked by the locations, a pair of items was directly associated with each other in the pair-association task. In this item-item association memory paradigm, the memory retrieval signal representing a target item reportedly appeared first in the perirhinal cortex of the MTL and spread backward to the visual area TE (29, 32, 33). Future studies should aim to determine whether the location signal in the HPC is retrieved from the item-cue information within the HPC (34) or if it is derived from other areas such as the perirhinal cortex (S5 Fig), which has been considered as a core brain region responsible for semantic dementia (35, 36) as well as a hub of converging sensory inputs (32, 37).

A generation of the retrieved location signal triggered by an item-cue was temporally separated from the perceptual signal input of a background-cue in the present study. The sequential presentations of the two cues allowed us to observe neuronal dynamics which may underlie the constructive process fitting the retrieved location information to the background-cue (S5 Fig). Sequential presentations of two cue stimuli were also applied in previous studies to investigate a conjunctive representation of the two stimuli in the HPC of rats (13) and in the perirhinal cortex of non-human primates (38). In these studies, animals learned to associate two- cues combination patterns (e.g., two sounds × two odors) directly with choice response (13) or reward delivery (38) because the combinations of two cue stimuli were arbitrarily assigned to their outcomes, which contrasts to the present study in which the combinations of the item- and background-cues necessarily determined the correct target positions (Fig 1c, see also S1 Text and S6 Fig). In contrast to the present study, memory retrieval signal appeared only after the second cue in both of the previous studies (13, 38). Interestingly, Terada et al. found that the two cue presentations (past and current events) and the expected choice (future event) were represented in the theta phase precession manner during the second cue period (13). Future studies should aim to determine whether the construction process shown in the present study is operated on the representational space defined by the phase precession.

Examination of the relationship between the retrieved location and the target location (Fig 4) revealed the presence of the default position/orientation for the background image in the HPC. Considering the training history of the monkeys for the item-location association, the default position may depend on their previous experiences. This finding may explain our mental representations of landmarks for their locations, which depends on our experiences (e.g., the Statue of Liberty in the New York City, bottom) (39). An unanswered question here is whether these mental locations were coded in the allocentric coordinates relative to the external world such as a frame of computer screen (20, 40), an egocentric location relative to the animals’ head position (41), or both (42).

Another unanswered question is whether the target-location-selective activity in the HPC encodes an action plan (i.e., endpoint of the saccade) or a mental representation of a target location. In the contextual fear memory paradigm (43), the HPC provides the amygdala with context information rather than its associated valence triggering fear responses (44). Moreover, recent human fMRI studies reported activation of the default mode network during future simulations (45), and suggested that the retrieved information spreads from the MTL to medial prefrontal cortex (PFC) (46), which is reportedly involved in decision-making (47). On the basis of these findings, we hypothesize that the HPC may provide its downstream regions with a target location, which may guide subsequent action selection (Fig 6) (48).

In the present CMP task, we found the three neuronal operations (i.e., *convergence, transference*, and *targeting*) in the HPC that were involved in the construction of goal-directed information. Constructive information processing is best recognized in the visual system (49). Based on the anatomical hierarchy, the construction proceeds from the retina to HPC through a large number of distinct brain areas to construct a mental image of an entire visual scene from local visual features (e.g., light spots, oriented bars). A recent electrophysiological study demonstrated constructive perceptual processing in the MTL, which combines an object identity with its location when the monkeys look at the visual object (20). However, a constructive process for perceiving an entire scene is still an unsolved question. As to the memory system, Schacter and his colleagues proposed the “*constructive episodic simulation hypothesis*” (1, 50), which assumes that our brain recombines distributed memory elements to construct either past episodes or future scenarios (i.e., “mental time travel”) (24, 25). However, the neuronal correlates to the constructive memory process for the mental time travel have not been identified. In the present study, we indicated that HPC neurons combined the memory and perception for flexible use of memory, which is in contrast to procedural or fear-conditioned memory (3-5). Considering its functional significance as declarative memory, this constructive process operated by the three neuronal operations in the HPC may be shared across species and might be a precedent of the constructive memory process combining only memory elements for the “mental time travel” in the evolution process of declarative memory system. The transitions of the three sequential neuronal operations for the construction process might be related with attractor dynamics substantiated by the HPC recurrent networks, which is reportedly involved in spatial memory of rodents (51-54). The underlying neuronal mechanisms would be investigated by theoretical and experimental study across species including both primates and rodents.

## Materials and Methods

### Experimental Design

#### Subjects

The subjects were two adult male rhesus monkeys (*Macaca mulatta*; 6.0–9.0 kg). All procedures and treatments were performed in accordance with the NIH Guide for the Care and Use of Laboratory Animals and were approved by the Institutional Animal Care and Use Committee (IACUC) of Peking University.

#### Behavioural task

We trained two monkeys on a constructive memory-perception (CMP) task (Fig 1). During both training and recording sessions, animals performed the task under dim light. The task was initiated by the animal fixating on a white square (0.4° visual angle) in the center of a display for 0.5 s. Eye position was monitored by an infrared digital camera with a 120-Hz sampling frequency (ETL-200, ISCAN). Then, an item-cue (diameter, 3.4°) and background-cue (diameter, 28.5°) were sequentially presented for 0.3 s each with a 0.7-s interval. After an additional 0.7-s delay interval, four equally-spaced white squares (0.4°) were presented at the same distance from the center (6°) as choice stimuli. One of the squares was a target, while the other three were distracters. The target was determined by a combination of the item-cue and the background-cue stimuli. The animals were required to saccade to one of the four squares within 0.5 s. If they made the correct choice, four to eight drops of water were given as a reward. When the animals failed to maintain their fixation (typically less than 2° from the center) before the presentation of choice stimuli, the trial was terminated without reward. Before the recording session, we trained the animals to associate two sets of four visual stimuli (item-cues) with four particular locations relative to the background-cue image that was presented on the tilt with an orientation from −90° to 90°. In order to avoid that the monkeys learn to associate each combination of the item-cue and the background-cue with a particular target location, the orientation of background image was randomized at a step of 0.1°, which increased the number of combinations (8 × 1800) and would make it difficult for the animals to learn all the associations among item-cues, background-cues and target locations directly. During the recording session, the item-cue was pseudorandomly chosen from the eight well-learned visual items, and orientation of the background-cue was pseudorandomly chosen from among five orientations (−90°, −45°, 0°, 45°, and 90°) in each trial, resulting in 40 (8 × 5) different configuration patterns. We trained the two monkeys using same stimuli but different item-location association patterns. The two monkeys performed the task correctly (chance level = 25%) at rates of 80.9% ± 8.1% (mean ± standard deviation; monkey B, n = 179 recording sessions) and 96.8% ± 3.1% (monkey C, n = 158 recording sessions) (S1 Fig).

#### Electrophysiological recording

Following initial behavioural training, animals were implanted with a head post and recording chamber under aseptic conditions using isoflurane anaesthesia. To record single-unit activity, we used a 16-channel vector array microprobe (V1 × 16-Edge; NeuroNexus) or a single-wire tungsten microelectrode (Alpha Omega), which was advanced into the brain by using a hydraulic Microdrive (MO-97A; Narishige) (11). The microelectrode was inserted through a stainless steel guide tube positioned in a customized grid system on the recording chamber. Neuronal signals for single units were collected (low-pass, 6 kHz; high-pass, 200 Hz) and digitized (40 kHz) (AlphaLab SnR Stimulation and Recording System, Alpha Omega). We made no attempt to pre-screen isolated neurons. Instead, once we succeeded in isolating any neuron online, we started a new recording session. The offline isolation of single units was performed using Offline Sorter (Plexon) by manual curation to make sure that noise transients were not included as units and that the same cell was not split into several clusters. The cells were isolated depending on the properties of spike waveforms. The cells were included into the analysis if the cells fired throughout the recording session with well-defined fields and a minimal mean firing rate as 1 Hz. On average, 128 trials were tested for each neuron (n = 456). The placement of microelectrodes into target areas was guided by individual brain atlases from MRI scans (3T, Siemens). We also constructed individual brain atlases based on the electrophysiological properties around the tip of the electrode (e.g., grey matter, white matter, sulcus, lateral ventricle, and bottom of the brain). The recording sites were estimated by combining the individual MRI atlases and physiological atlases (55).

The recording sites covered between 3 and 16 mm anterior to the interaural line (monkey B, left hemisphere; monkey C, right hemisphere; S2 Fig). The recording sites cover all the subdivisions of the HPC (i.e., dentate gyrus, CA3, CA1, and subicular complex) (11). A final determination will require future histological verification (both animals are currently still being used).

### Statistical Analysis

All neuronal data were analysed by using MATLAB (MathWorks) with custom written programs, including the statistics toolbox.

#### Classification of task-related neurons

For the item-cue period, we calculated mean firing rates of eight consecutive 300-ms time-bins moving in 100-ms steps, covering from 0- to 1000-ms after item-cue onset, across all correct trials. We evaluated the effects of “item” for each neuron by using one-way ANOVA with the eight item-cue stimuli as a main factor (*P* < 0.01, *Bonferroni correction* for eight analysis-time windows). We referred to neurons with significant item effects during any of the eight analysis-time windows as item-selective neurons. For the background-cue period, we calculated the mean firing rates of eight consecutive 300-ms time- bins moving in 100-ms steps, covering from 0- to 1000-ms after the background-cue onset. We evaluated the effects of “co-location,” “background,” and “target” for each neuron by using three-way nested ANOVA with the four co-locations, three background-cue orientations, and four target locations as main factors, and the eight item-cues nested under the co-locations (*P* < 0.01, *Bonferroni correction* for eight analysis-time windows). The three-way nested ANOVA was conducted using the correct trials with −90°, 0°, and 90° background-cues.

#### Analysis of retrieval signal during item-cue period

To show the time course of activity for individual item-selective neuron, a spike density function (SDF) was calculated using only correct trials and was smoothed using a Gaussian kernel with a sigma of 20 ms. To examine the retrieval signal, we assessed the relationship between the responses of co-location items by using Pearson correlation. For each item-selective neuron, we calculated mean firing rates for each 300-ms time-bin moving by 100 ms during the item-cue period for each correct trial (8 bins in total). Then, we performed one-way ANOVA and calculated a grand mean to each item stimulus across correct trials for each bin. For the bins that showed a significant (*P* < 0.01, *Bonferroni correction* for eight analysis-time windows) item effect, we calculated the correlation coefficients between the mean firing rates of co-location items. We then averaged *Z*-transformed values of the correlation coefficients across time-bins for each neuron. The average value was finally transformed into *r* values (i.e., co-location index) as shown in Figure 2.

#### Analysis of task-related signal during background-cue period

To show the time course of activity for individual task-related neuron, a SDF was calculated using only correct trials and was smoothed using a Gaussian kernel with a sigma of 20 ms. For comparing time courses of proportions of task-related (co-location, background, and target) neurons and their signal amplitudes, we conducted three-way nested ANOVA for each 100-ms time-bin moving by 1 ms to test significances (*P* < 0.01, uncorrected) with *F* values for each neuron. The three-way nested ANOVA was conducted using only the correct trials with −90°, 0° and 90° background-cues.

#### Analysis of similarity of orientation tuning

To evaluate the effects of background-cue on the co-location-selective responses, we used data in the correct trials with −90°, 0°, and 90° background-cues. We first calculated mean firing rates for each co-location during the 60–1000- ms period from an onset of the background-cue and determined the “best co-location” and the “second-best co-location” based on the mean firing rates for each neuron. We then calculated Pearson correlation between responses to the different orientations of background-cues for the best co-location and those for the second-best co-location (i.e. similarity of tuning). We also examined a time course of the background-cue effect on the co-location-selective responses by calculating the population-averaged correlation coefficients for each 100-ms time-bin moving by 1 ms.

#### Calculation of matching index

To evaluate the relationship between the retrieved location and the target location, we used data in the correct trials with −90°, 0°, and 90° background-cues. For each neuron exhibiting both item-selectivity during item-cue period and target-selectivity during background-cue period, we first averaged responses during the 60- to 1000-ms period from item-cue onset in each trial and calculated a grand mean across trials to each of the four co-locations. In addition, we averaged responses during the 60- to 1000-ms period from background-cue onset in each trial and calculated a grand mean across trials to each of the four target locations. According to the three potential matching patterns, we sorted the firing rates to the co-locations and calculated Person correlation coefficients between responses to the co-locations in each of the three potential matching patterns and those to the target locations

#### Analysis of target signal

To evaluate background-cue effect on the target signal, population-averaged SDFs (best – other target locations) were calculated for target-selective neurons across the correct trials with −90°, 0°, and 90° background-cues. We first averaged responses during the 60- to 1000-ms period from background-cue onset in each trial and calculated a grand mean across correct trials to each of the four target locations to determine the “best target location” for each neuron. The SDFs to each orientation (−90°, 0°, and 90°) of background-cues for all target locations were normalized to the amplitude of the mean response to the best target location, and the normalized SDFs for the best target location was subtracted by the mean normalized responses across the other target locations. The population-averaged SDFs (i.e., target-selective response) were smoothed using a Gaussian kernel with a sigma of 20 ms.

#### Error analysis

We examined whether the target-selective activities signaled positions the subjects chose or correct positions during the background-cue period in error trials by using a partial correlation coefficient. To calculate the partial correlation coefficient for each neuron, we first calculated an average firing rate during each 300-ms time-bin moving by 100 ms during the background-cue period (i.e., eight time-bins in total) for each target position (i.e., eight positions in total) across the correct trials. We next prepared for three arrays for each neuron containing “*n*” elements in each array (“*n”* is the number of error trials for each neuron): (i) firing rates in the *i-th* error trial (*i* = 1 to *n*) (dependent variable, ***D***); (ii) the mean firing rate across correct trials with the same target position as the subject chose in the *i-th* error trial (explanatory variable, ***X***); (iii) the mean firing rate across correct trials with the same target position as the subject missed (i.e., correct answer) in the *i-th* error trial (explanatory variable, ***Y***). The partial correlation coefficients of the dependent variable, ***D*** with explanatory variables, ***X*** and ***Y*** were calculated in each time-bin for each neuron when the neuron’s responses in correct trials showed a significant target effect (*P* < 0.01, *Bonferroni correction* for eight analysis-time windows) and the mean firing rate across trials was larger than 1 Hz in that time-bin. The mean partial correlation coefficients were calculated across the active time bins (i.e., *P* < 0.01, *Bonferroni correction* for eight analysis-time windows, >1 Hz) for each neuron using *Z*-transformation.

## Acknowledgments

We thank S. Kitazawa, K.W. Koyano, W.A. Suzuki, I. Lee, K. Miyamoto, S. Fujisawa, H. Chen, H. Deng for helpful comments and S. Xue for expert animal care. We thank J. Gao, W. Men, G. Yang, and the National Center for Protein Sciences at Peking University for assistance with MRI
scanning.

## Supporting Information

### Supporting text

**S1 Text. Training procedures for the CMP task** In order to prevent the monkeys from learning to associate each combination of the item-cue and the background-cue with a particular target location through repetitive trials by positive reinforcement learning, we first trained the monkeys to learn the task rule of the CMP task using a training stimulus set in the preliminary training before the final training using a recording stimulus set. For the training stimulus set, we used monochromatic simple-shaped objects (e.g., cross, heart) as item stimuli and a large disk with four monochrome colors in individual quadrants as the background stimulus (S6 Fig). The monkeys were required to touch the target location on the touch screen (3M^TM^ MicroTouch^TM^ Display M1700SS) according to the combination of item-cue and background-cue without any fixation requirement during the task. We used a fixed orientation (0°) of the background image as the background-cue during the initial period of the preliminary training. After the animals learned the combination between items and the locations on the 0° background, we presented the oriented-background image as background-cue. The orientation of the background image was randomly chosen from −90° to 90° (0.1° step). After they learned the CMP task using the training stimulus set, we trained them to learn the stimulus set for the recording (Fig 1a). We found the animals acquired the final stimulus set faster than the training set, which may imply they learned the task rule during the preliminary training and applied it to the recording stimulus set. Finally, we trained the monkeys to answer the target location by saccade. Once the monkeys reached the criterion of 70% accuracy, we started the recording session.

**S2 Text. Examinations of neuronal responses to co-location stimuli and eye positions** We found that item-selective neurons showed similar amplitudes of activities to item stimuli that were assigned to the same location relative to the background-cue stimulus, implying that the item-selective responses reflected to-be-retrieved location information. One potential caveat to this interpretation would be that the location signal of the item-selective neurons reflected animals’ eye positions rather than the item-location association memory. To examine this possibility, we first evaluated the co-location effects of item-cue stimuli on eye positions during the item-cue period when we collected spike-firing data of item-selective neurons. We grouped trials with two different item-cues sharing the same location into one type of co-location trials. We conducted one-way ANOVA with four types of co-location trials as the main factor for the recording session of each item-selective neuron, and found that 37% of recording sessions showed significant (*P* < 0.025, either horizontal or vertical) effects of co-location trials on eye position (item-selective neurons with recorded eye data, n = 89). Then we separated item-selective neurons into two groups according to the co-location effects of item-cue stimuli on eye positions during the recording sessions. If the location signal of the item-selective neuron was explained by the eye positions, the location signal should be stronger when the co-location of item-cue affects eye positions more. We examined whether or not the item-selective responses signaled the location information during the item-cue period by calculating the correlation coefficients between the responses to co-location stimuli. The correlation coefficients were even larger for the item-selective neurons sampled during the recording session without significant co-location effects on the eye positions (*r* = 0.82, median) than those with significant effects (*r* = 0.73). These results indicate that the location information exhibited by item-selective neurons could not be explained by the animals’ eye positions. Because of hardware defects, we failed to record eye positions during the early period of the experiment (47 of 136 item-selective neurons).

**S3 Text. Detection of task-related signals during the background-cue period** To quantify neuronal responses during the background-cue period, we performed three-way nested ANOVA with co-location, background-cue, and target as the main factors for each neuron. Item-cues were nested under the co-location factor, and their effects were negligible during the background-cue period in all areas. In this analysis, we used data only in correct trials whose background-cues were −90°, 0°, and 90° because −45° and 45° background-cues bring about target positions (i.e., top, right, bottom, and left) that were different from those of the −90°, 0°, and 90° background- cues (top-right, bottom-right, bottom-left, and top-left). Because of the limited orientations of the background-cue, a co-location would theoretically bring about a bias for a target location (e.g., co-location I could result in top-right, bottom-right and top-left, but not bottom-left target location). However, we did not find an effect of this confounding factor on the detection of target-selective neurons at least before background-cue presentation because we found only a negligible number (around the 1% of chance level) of target-selective neurons, while a large proportion (16%, S1 Table) of target-selective neurons were noted after background-cue presentation.

### Supporting figures

**S1 Fig.**
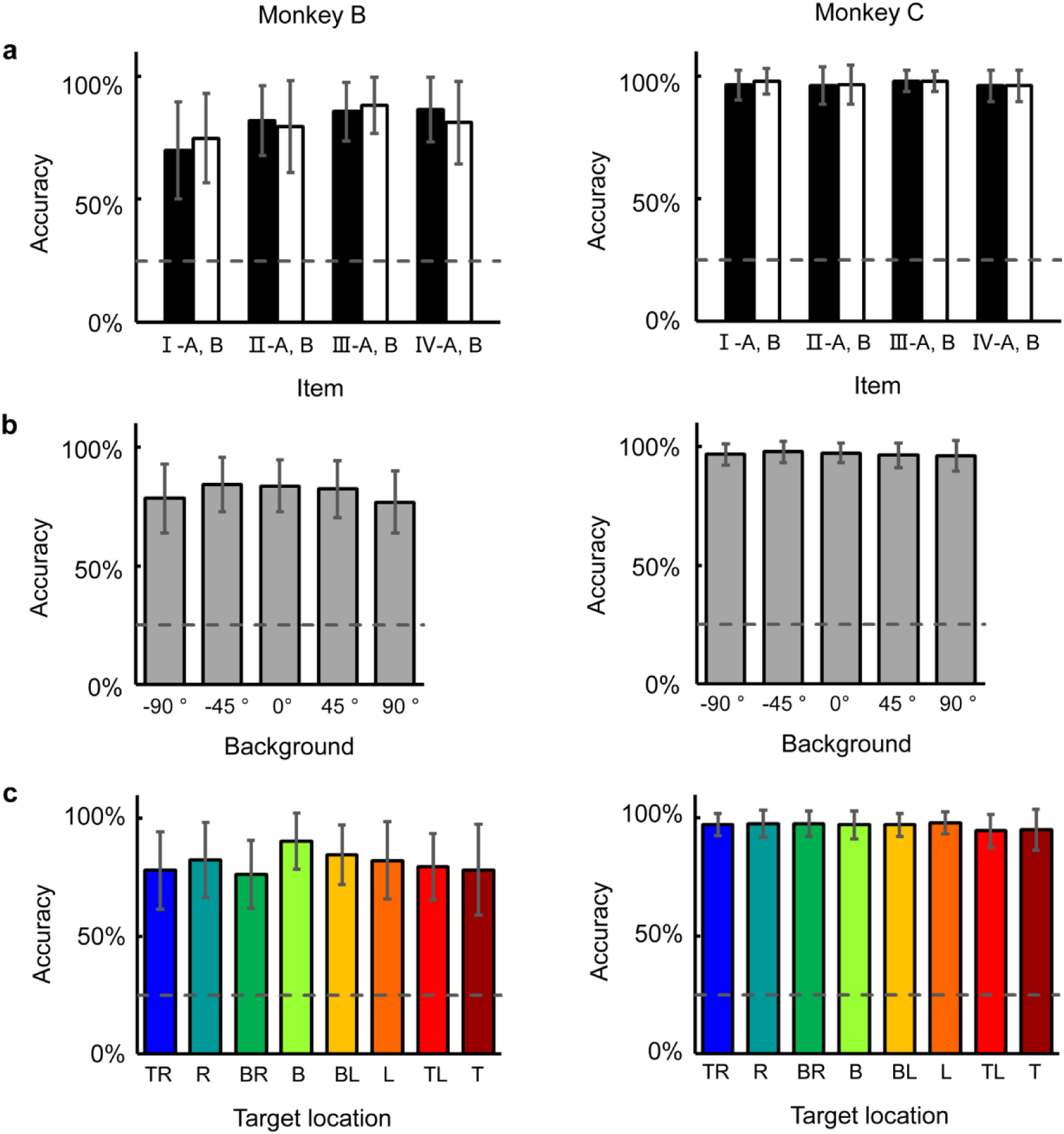
Performance in the CMP task. Performance of each monkey in task-related conditions. Monkey B, n = 179 recording sessions. Monkey C, n = 158 recording sessions). Error bar, standard deviation. Dashed line, chance level = 25%. **(a)** Performance for 8 item stimuli as item-cue. Black bars, set A. White bars, set B. **(b)** Performance for 5 orientations of background-cue. **(c)** Performance for 8 positions on the display as target locations. TR, top-right. R, right. BR, bottom-right. B, bottom. BL, bottom-left. L, left. TL, top-left. T, top.

**S2 Fig.**
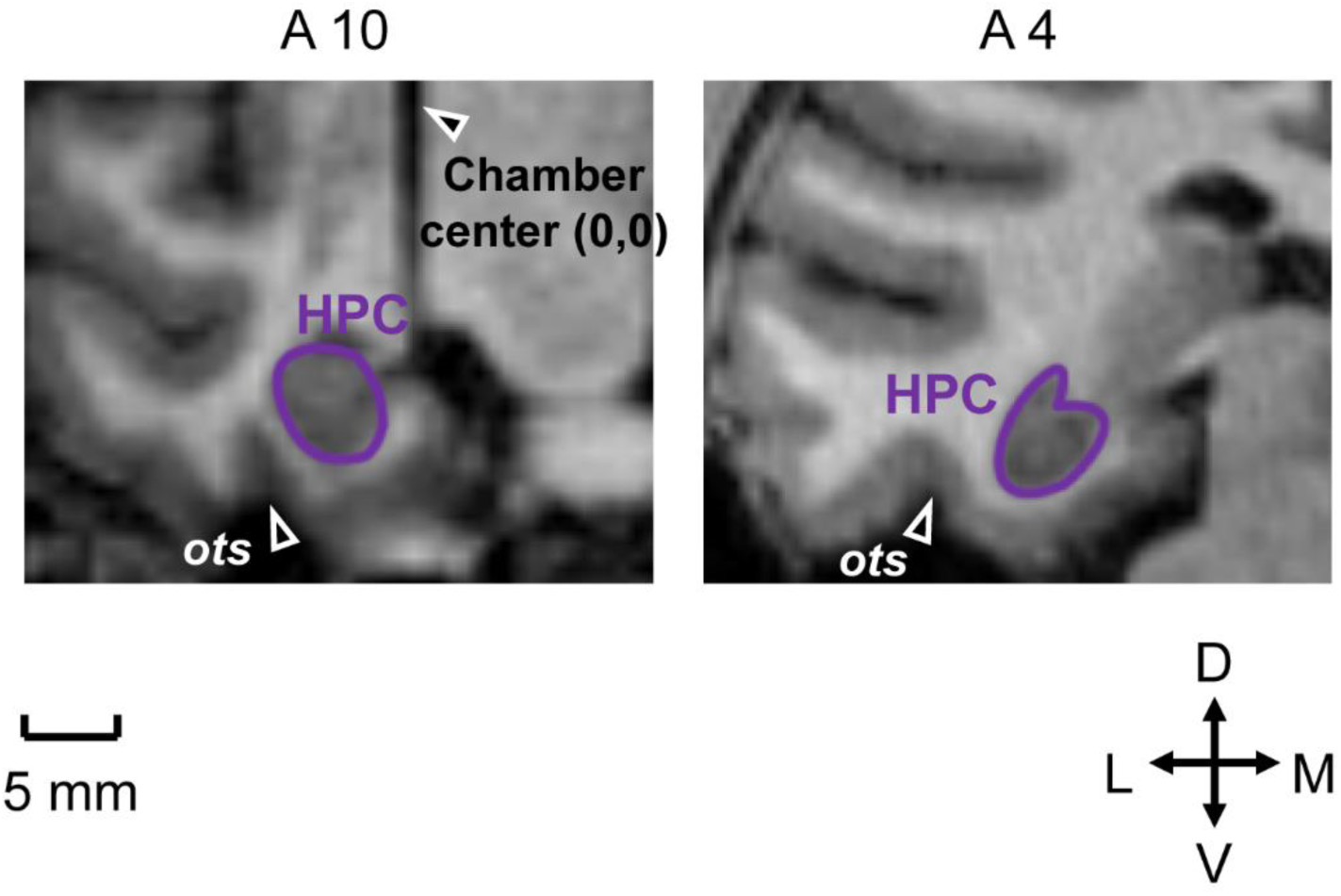
Recording region. MRI images corresponding to the coronal planes anterior 4, 10 mm from interaural line of monkey C (right hemisphere). Recording region is the hippocampus (HPC). A reference electrode implanted in the center of chamber was observed as a vertical line of shadow in the coronal plane at A10. *ots*, occipital temporal sulcus. D, dorsal. V, ventral. L, lateral. M, medial.

**S3 Fig.**
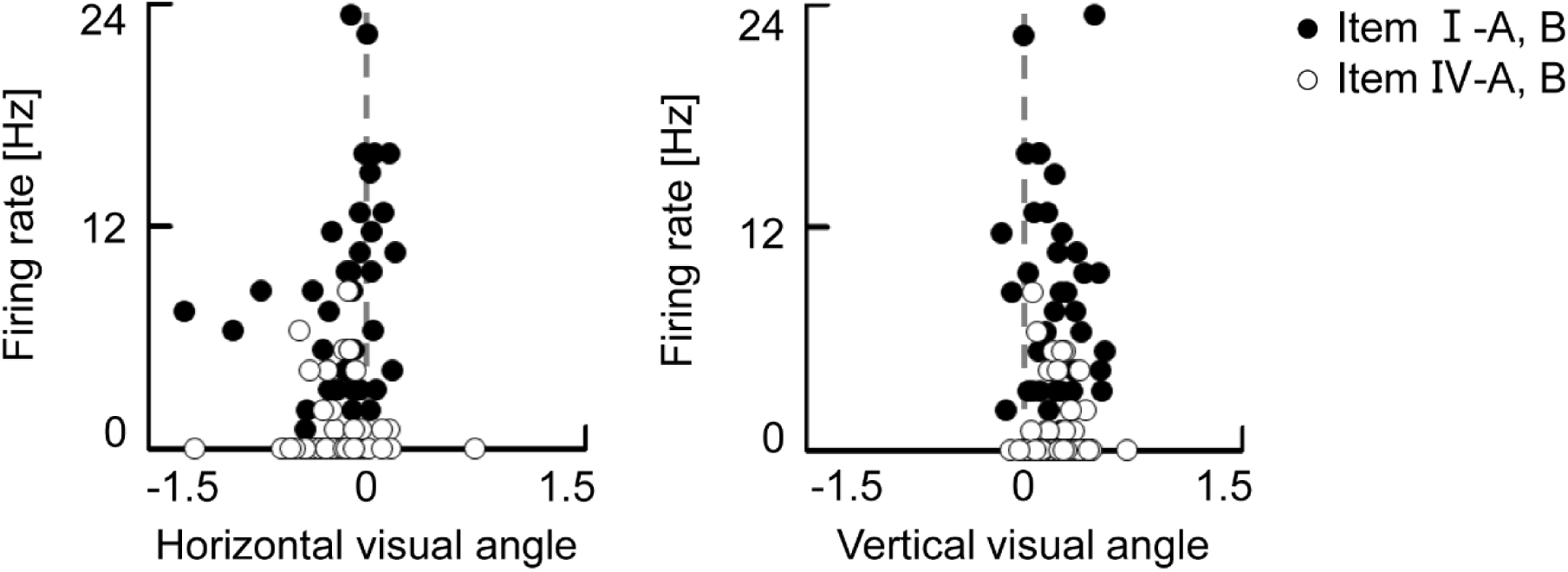
Examinations of neuronal responses and eye positions. Firing rates plotted as a function of eye positions during the item-cue period for the neuron shown in Figures 2a and 2b. Each dot indicates one trial. Black dots indicate trials with the best co-location stimuli as item- cues. White dots indicate trials with the worst co-location stimuli as item-cues. The large overlaps were found in the distributions of the eye positions between the trials with the best and worst co-location items (*P* = 0.16 for horizontal, *P* = 0.25 for vertical, *t*-test, two-tailed) while distributions of the firing rates were significantly different between the two trial types (*P* < 0.0001). These results indicate that the item-selective responses shown in Figures 2 cannot be explained by the animal’s eye positions.

**S4 Fig.**
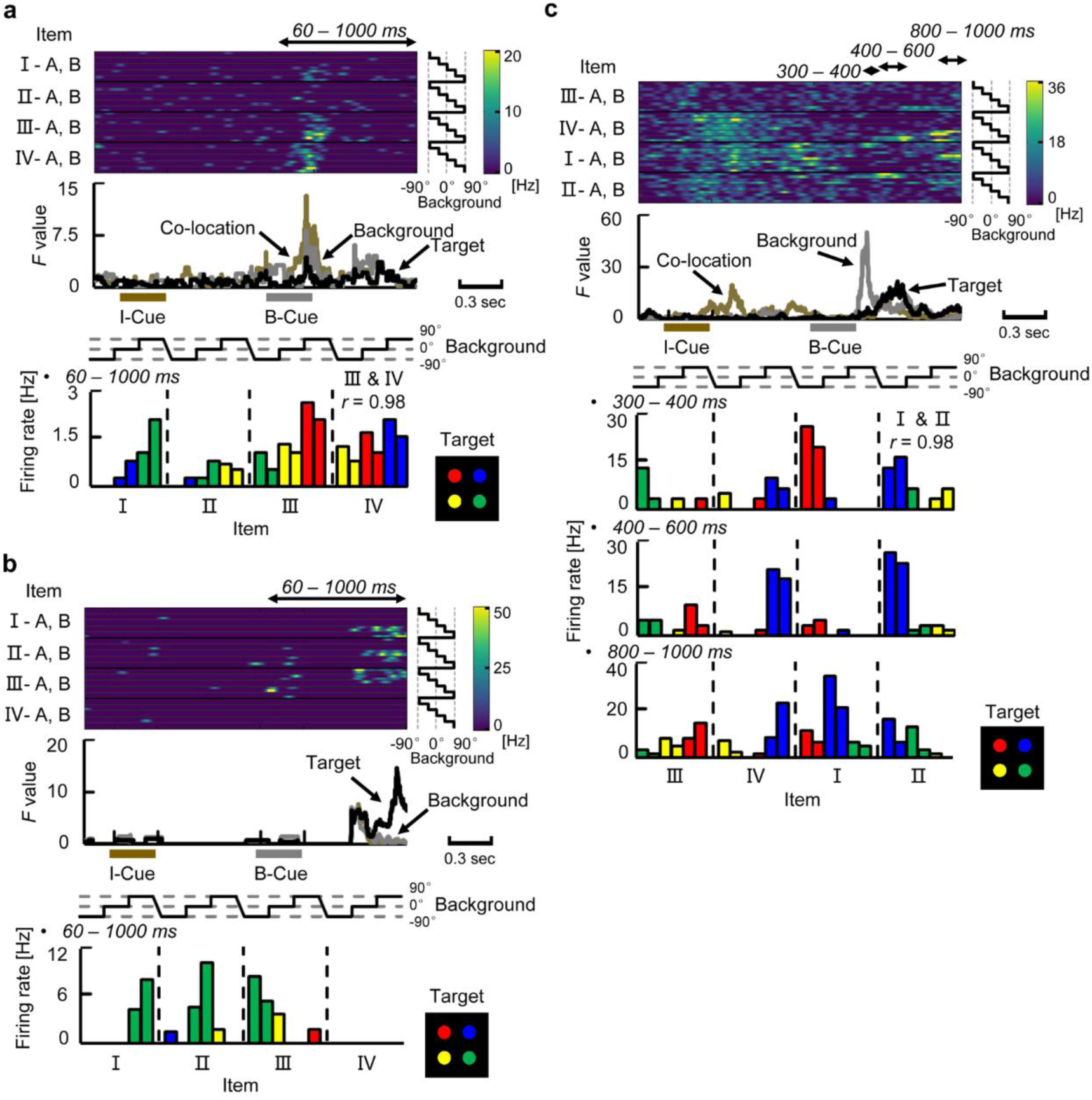
Example neurons involved in the constructive process. **(a)** Example neuron signaling co-location and background-cue in a “*convergent*” manner. This neuron did not show item-cue selective responses during the item-cue period (*P* = 0.34, one-way ANOVA), but it exhibited the co-location-selective responses during the background-cue period (*P* < 0.01, three-way nested ANOVA). The background-selective responses were combined with the co-location-selective responses. The preferred orientation of the background-cue stimulus was 90° for this neuron across co-locations. The same format as Fig 4a. **(b)** Example neuron signaling a “*targeting*” location. This neuron did not show item-selective responses during the item-cue period (*P* = 0.81, one-way ANOVA), but it exhibited the target-selective responses during the background- cue period (*P* < 0.0001, three-way nested ANOVA). The best target location was bottom-right of display (*green*). **(c)** Example neuron that changed the activity patterns, showing multiple operations for the construction (i.e. *convergence, transference*, and *targeting*) during the background-cue period. This neuron showed item-selective responses during the item-cue period (*P* < 0.0001), and the preferred items in the item-cue period were I-A, I-B, IV-A, and IV-B. During 300-400 ms after background-cue onset, the background-selective responses were “converged” on the co-location-selective responses and the preferred orientation was −90° across co-locations. During 400-600 ms after background-cue onset, this neuron exhibited strong responses only to the particular combinations of item-cue and background-cue “*transferring*” to the top-right target position (*blue disk*) (*blue bars*, II-A & II-B, −90° and IV-A & IV-B, 90°). During 800-1000 ms after background-cue onset, this neuron exhibited selective responses to all the combinations of item-cue and background-cue “*targeting*” for the top-right target position.

**S5 Fig.**
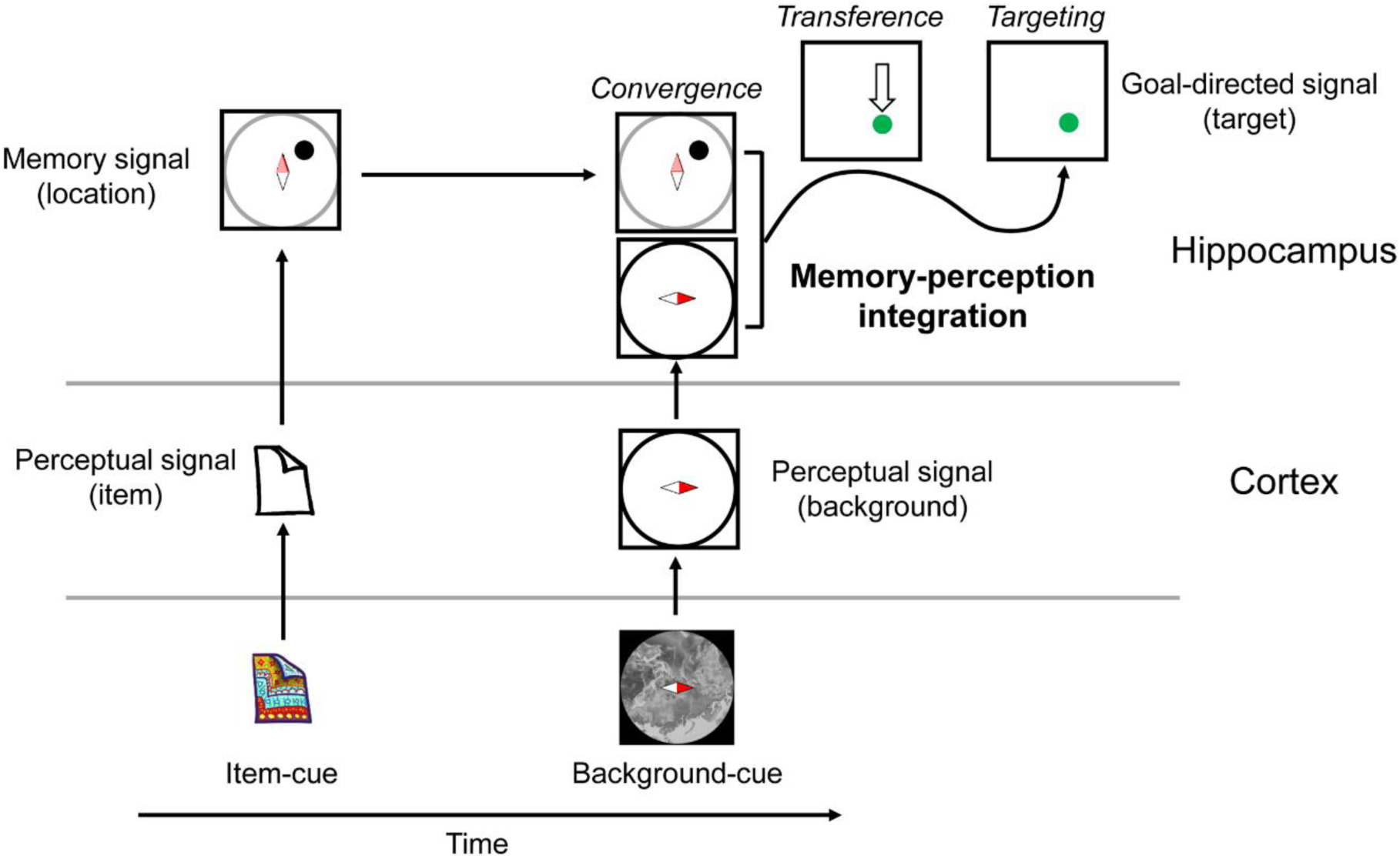
Constructive process for the flexible use of memory. Schematic diagram of neuronal signals during a trial of the CMP task, in which the item-cue and the orientation of background- cue were I-B and 90°, respectively. In the HPC, the retrieved location of the item is represented relative to the 0° background image. The incoming perceptual signal is integrated with the memory signal to construct an updated information signaling the target location by following sequential neuronal operations: *convergence* [i.e., memory (co-location I on the 0° background) + perception (90° background)], *transference* [i.e., from the top-right on the display (co-location I on the 0° background) into the bottom-right on the display (co-location I on the 90° background)], and *targeting* (i.e., coding bottom-right on the display).

**S6 Fig.**
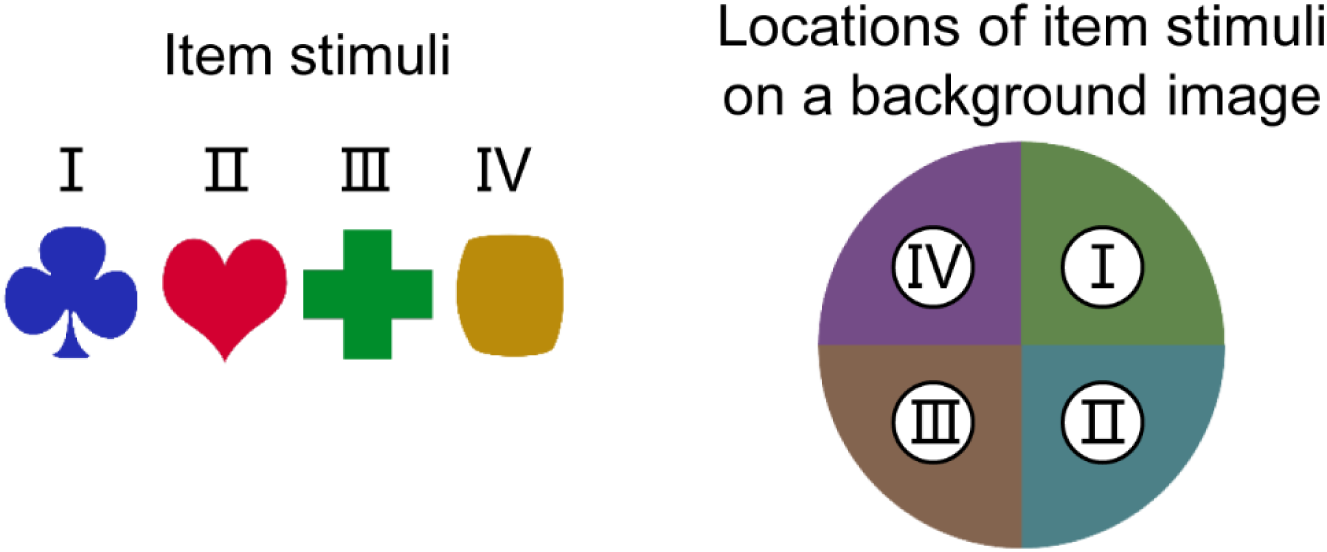
Stimuli for preliminary training of the CMP task. (Left) Simple shape objects with monochrome colour as item stimuli. (Right) Large disk with four monochrome colours in individual quadrants as a background stimulus. Each item stimulus was assigned to one location on the background stimulus.

### Supporting table

**S1 Table.**
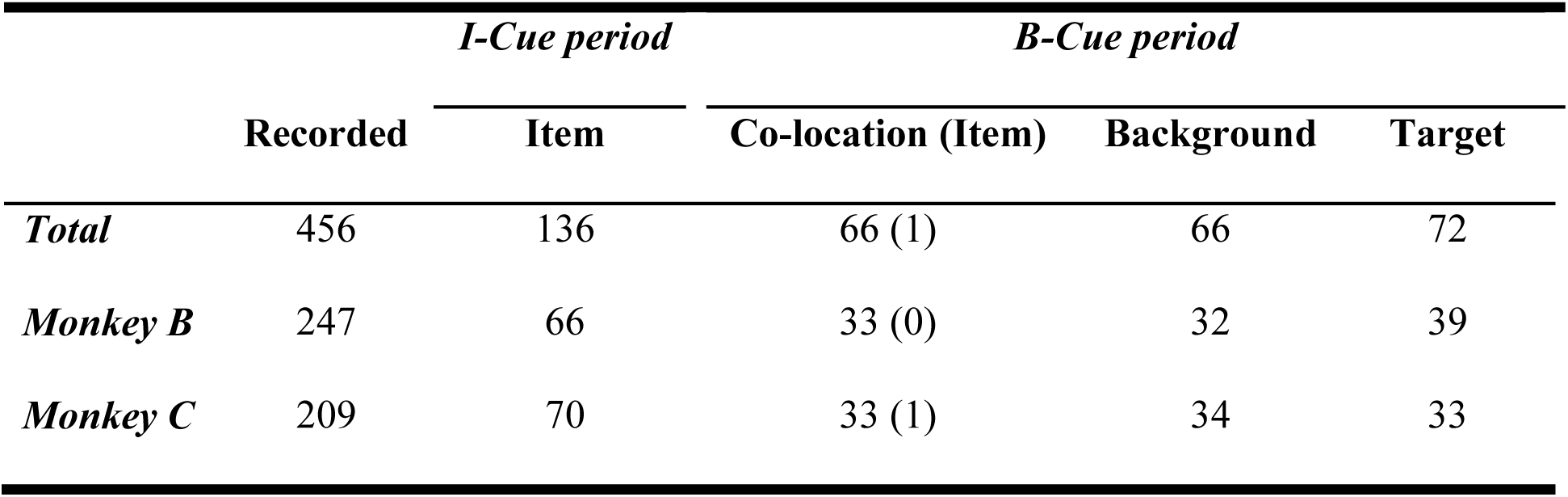
Numbers of task-related neurons. Numbers of neurons showing item effect during the item-cue period, and those showing item, co-location, background, and target effects during the background-cue period at a significance level of *P* < 0.01. “Item” indicates an item effect during the item-cue period or the background-cue period. “Co-location” indicates a co-location effect during the background-cue period. “Background” indicates a background effect during the background-cue period. “Target” indicates a target location effect during the background-cue period.

